# Mapping the genetic landscape of biomineralization in *Magnetospirillum magneticum* AMB-1 with RB-Tnseq

**DOI:** 10.1101/2021.08.13.456315

**Authors:** Hayley C. McCausland, Kelly M. Wetmore, Adam P. Arkin, Arash Komeili

## Abstract

Magnetotactic bacteria (MTB) are a phylogenetically diverse group of bacteria remarkable for their ability to biomineralize magnetite (Fe_3_O_4_) or greigite (Fe_3_S_4_) in organelles called magnetosomes. The majority of genes required for magnetosome formation are encoded by a magnetosome gene island (MAI). Most previous genetic studies in MTB have focused on the MAI, using screens to identify key MAI genes or targeted genetics to isolate specific genes and their function in one specific growth condition. Here, we conducted random barcoded transposon mutagenesis (RB-TnSeq) in *Magnetospirillum magneticum* AMB-1 to identify the global genetic requirements for magnetosome formation under different growth conditions. We generated a library of 184,710 unique strains in a wild-type background, generating ~34 mutant strains for each gene. RB-TnSeq also allowed us to determine the essential gene set of AMB-1 under standard laboratory growth conditions. To pinpoint novel genes that are important for magnetosome formation, we subjected the library to magnetic selection screens in varied growth conditions. We compared biomineralization in standard growth conditions to biomineralization in high iron and anaerobic conditions, respectively. Strains with transposon insertions in the MAI gene *mamT* had an exacerbated biomineralization defect under both high iron and anerobic conditions compared to standard conditions, adding to our knowledge of the role of MamT in magnetosome formation. Mutants in *amb4151*, a gene outside of the MAI, are more magnetic than wild-type cells under anaerobic conditions. All three of these phenotypes were validated by creating a markerless deletion strain of the gene and evaluating with TEM imaging. Overall, our results indicate that growth conditions affect which genes are required for biomineralization and that some MAI genes may have more nuanced functions than was previously understood.

## Introduction

Magnetotactic bacteria (MTB) are a diverse group of bacteria capable of producing intracellular organelles called magnetosomes [1–7] after taking up iron from the surrounding environment. Magnetosomes are membrane-bound compartments in which biomineralization of magnetic crystals of magnetite (Fe_3_O_4_) and/or greigite (Fe_3_S_4_) occurs [4]. Crystals are organized into linear chains along the long axis of the cell, forming a magnetic dipole that allows the cell to orient towards Earth’s magnetic fields [8]. MTB inhabit low-oxygen environments and are typically found at the oxic-anoxic transition zone (OATZ) in a water column [4]. Navigation along magnetic field lines is thought to allow cells to efficiently locate the OATZ; a process called magnetoaerotaxis [9].

MTB are ubiquitous in aquatic environments and can account for up to 30% of microbial biomass in some habitats [5]. Because of their presence in water and the large amounts of iron that each cell captures in the process of biomineralization, it is likely that MTB have a large impact on iron cycling in the ocean, potentially taking up anywhere from 1-50% of dissolved iron inputs into the ocean [10]. However, most of what we know about MTB in the environment comes from surveys of species and their numbers in particular habitats. The dynamic responses of MTB, particularly at a genetic level, to fluctuations in the environment remain largely unexplored. A greater understanding of the molecular mechanisms of biomineralization and iron sequestration by MTB in response to changing conditions will inform environmental studies, including the impact that MTB have on iron cycling. Yet, most of our knowledge of the genetics and cell biology of MTB comes from studies done under static conditions that do not accurately reflect the natural environment of MTB grow. Here, we take advantage of the well-studied model organism *Magnetospirillum magneticum* AMB-1 (AMB-1) and develop a high throughput genetic strategy to connect genetics to environmental changes.

The genes needed to produce magnetosomes are encoded in magnetosome gene clusters (MGCs). In some species—like the model organisms AMB-1 and *Magnetospirillum gryphiswaldense* MSR-1 (MSR-1)—the MGCs are discreet magnetosome gene islands (MAIs). The MAIs in AMB-1 and MSR-1 are well characterized [11,12]. Both organisms have approximately 100 genes in their respective MAIs. However, it was shown by Kolinko et al. that only 31 of the genes in the MSR-1 MAI are necessary and sufficient to synthesize magnetosomes—though even fewer may be required [13]. This implies that the other ~70 genes in the MAI are either unnecessary for the formation of magnetosomes, are redundant with other genes, or are only required under certain growth conditions. In fact, some MAI genes are only conditionally required for magnetosome formation. For example, the genes *mamX* and *ftsZm* are necessary under oxygen-reducing conditions but not nitrate-reducing conditions [14,15].

Genes outside the MSR-1 MAI (ex-MAI genes) have also been connected to magnetosome formation as growth conditions are changed. Deletion of the *nap* operon in MSR-1 resulted in the formation of small, poorly aligned magnetosomes [16], indicating that nitrate metabolism, while not necessary for the growth of MSR-1, is critical in the formation of magnetosomes. The possibility that other genes—both inside and outside the respective MAIs of MSR-1 and AMB-1—are required for magnetosome formation under specific conditions has not been thoroughly examined [17–21].

Previous genetic studies in magnetotactic bacteria have used either targeted reverse genetics to determine the role of individual genes or operons, or forward genetic screens that focus solely on clear magnetosome mutants. Here, we present a global and readily scalable approach to investigate the genetic requirements of AMB-1. We used a screening technique called random barcoded transposon site sequencing (RB-TnSeq) [22], which involves generating a pooled library of tens of thousands of transposon mutants. First, we used the RB-TnSeq library to determine the essential gene set of AMB-1 in our standard growth conditions, providing a broader view of the lifestyle of this important model organism. Then, we used the RB-TnSeq library to conduct a high-throughput magnetic selection to study the genetic requirements for magnetosome formation under multiple environmental conditions. In particular, we focused on magnetosome formation in varying oxygen and iron concentrations, which have both been shown to influence magnetosome formation [23,24]. Magnetic selection experiments uncovered new functions of known magnetosome genes, while also revealing novel ex-MAI genes that may have a role in magnetosome formation.

## Results

### RB-TnSeq libraries reveal the essential gene set of AMB-1

To conduct high-throughput genetic screens in AMB-1, we generated pooled transposon insertion libraries containing thousands of mutants using the transposon mutagenesis technique, RB-TnSeq [22] (Fig.1). A library of ~30 million *mariner* transposon vectors (APA752) in the *E. coli* host strain WM3064 was conjugated with AMB-1. After growth on selective plates, all colonies were pooled for sequencing. The location of each transposon was mapped to the AMB-1 genome using Illumina sequencing.

**Figure 1.**
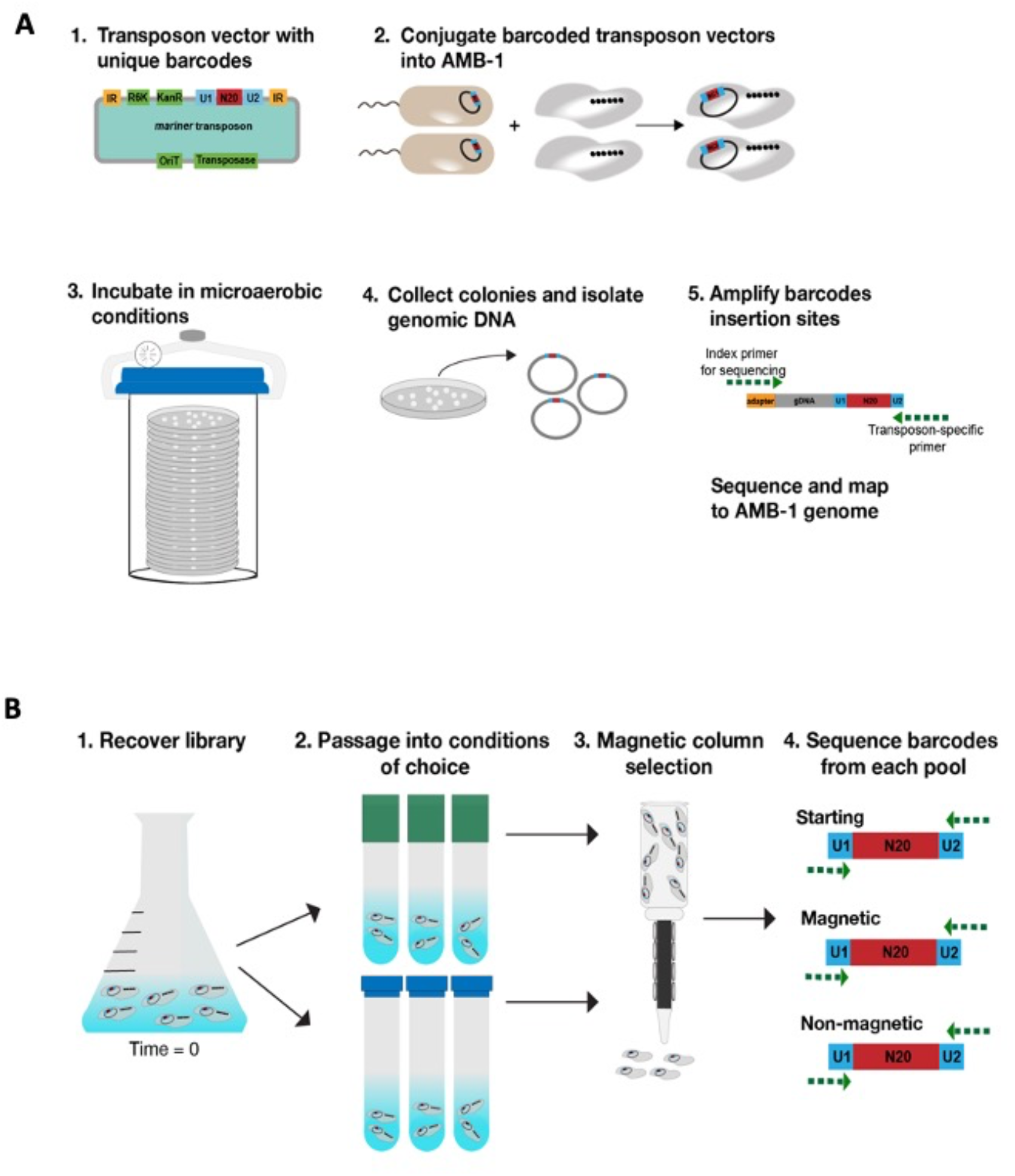
RB-TnSeq and magnetic selection. (A) Flow chart of RB-TnSeq library construction and sequencing. 1 (B) Flow chart of magnetic selection using RB-TnSeq library

We generated a successful library in WT AMB-1 called magnetotactic bacteria library ML2 (ML2). The library was constructed in our standard laboratory conditions: MG media supplemented with 30μM iron and grown microaerobically. ML2 contained 183,760 unique barcodes (Fig. 2A, B). There were 34.3 hits per protein on average (mean). The read bias (mean:median reads per hit protein) of 1.74 indicates a moderate bias in the library, where no genes are underrepresented by more than two-fold, on average. Since ML2 has a large number of mutant strains with broad coverage of the genome, it was used for downstream biomineralization screens.

**Figure 2.**
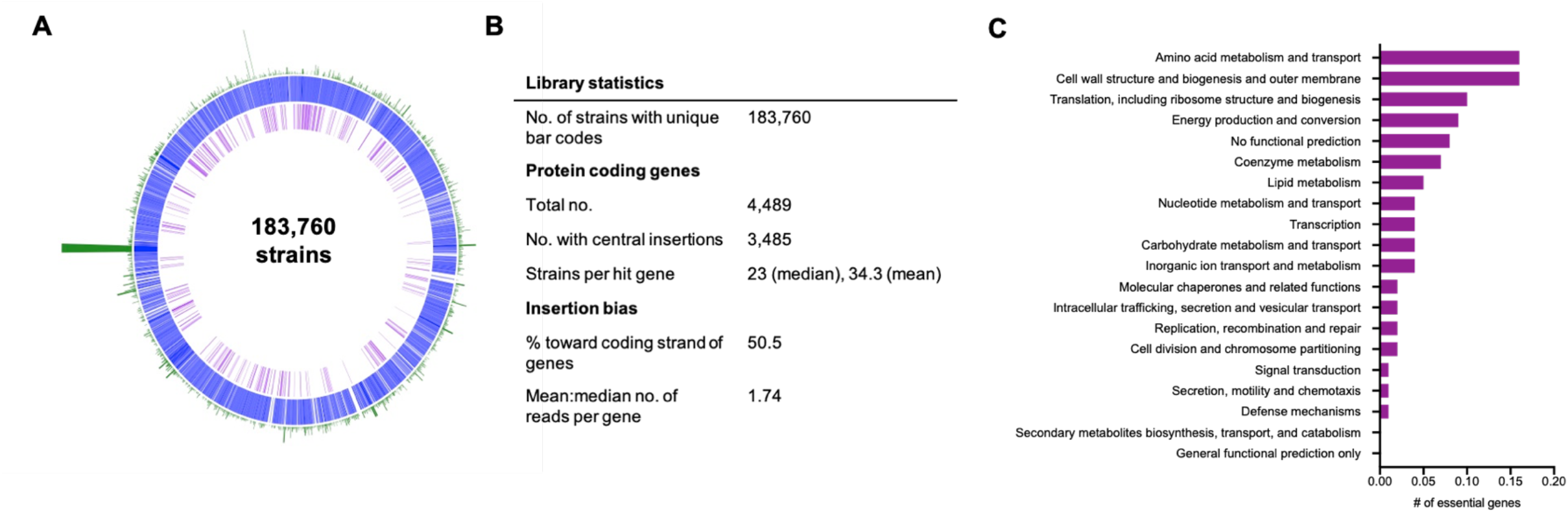
RB-TnSeq libraries and essential gene sets (A) Map of WT AMB-1 RB-TnSeq library showing AMB-1 genome (blue), WT essential genes (purple), and transposon insertion loci and densities (green) (B) Summary of library statistics for Magneto ML2. (C) Genes essential to the Magneto ML2 library under standard growth conditions grouped into COG categories

Another benefit of making an RB-TnSeq library is that it can be used to determine the putative essential gene set of a strain in the conditions under which it was made (Fig. 2A). Essential genes have very few or no insertions because those mutants should not be viable. Here, we determined the essential gene set of ML2 under standard laboratory conditions. We identified 445 essential genes in the AMB-1 genome—~9.9% of protein coding genes, which is similar to the percentage of essential genes in other RB-TnSeq libraries [25]. The remaining 4216 genes are either non-essential (3774 genes) or were not included in the list of essential genes (472 genes). Those genes that were not included are very similar to other regions of the genome or less than 800 nucleotides. Genes larger than 800 nucleotides were determined to be unlikely to have no insertions by chance [25].

Binning the essential genes into COG categories showed that amino acid metabolism and cell wall structure/biogenesis accounted for the largest proportion of essential genes (16% each) (Fig. 2C). Translation (10% of essential genes) and energy production (9% of essential genes) also contained a large proportion of essential genes. While these categories are expected for essential genes, the individual genes within each category can provide information about what MTB need to grow. For example, a succinate dehydrogenase subunit (*amb3952* or AMB_RS20000) and tartrate dehydrogenase (*amb3176* or AMB_RS16020) are essential under standard conditions, indicating the role of the TCA cycle in heterotrophic metabolism of AMB-1. Additionally, nitrate reductase subunits (*amb0531* and *amb0533* or AMB_RS02735 and AMB_RS02745), nitrate ABC transporter (*amb0534* or AMB_RS02750), and nitrate/sulfonate/bicarbonate ABC transporter ATP-binding protein (*amb0535* or AMB_RS02755) are essential, where nitrate was the only alternative electron acceptor available for respiration.

### Magnetic selection reveals genes important for biomineralization under high iron conditions

To screen for mutants with defects in biomineralization, we used a magnetic column to separate magnetic from non-magnetic cells in the RB-TnSeq library (Fig. 1B). After thawing an aliquot of ML2 and growing to late log phase (OD_400_ ~0.150), cells were passaged into growth conditions of choice and allowed to grow to stationary phase (OD_400_ ~0.250). Then, each culture was filtered through a magnetic column lined with magnets. Both the non-magnetic and magnetic samples were collected. A sample of the pre-column culture was saved as a control. BarSeq (sequencing of the unique barcodes) was performed on magnetic, non-magnetic, pre-column, and time zero samples and then a magnetic column score (MCS) for each gene was calculated based on each strain’s abundance (Fig. 3). It should be noted that BarSeq experiments are usually used to measure growth of each strain, based on a strain’s abundance in each experiment. Here, we have adapted the measure of strain fitness to evaluate biomineralization capabilities based on abundance of each strain in pre-column, magnetic, and non-magnetic samples. To account for any growth defects that might affect MCS, we normalized by subtracting pre-column gene MCS from magnetic and non-magnetic gene MCS before comparing to time zero. Raw fitness scores for all experiments can be viewed in the Fitness Browser (http://fit.genomics.lbl.gov) [25].

**Figure 3.**
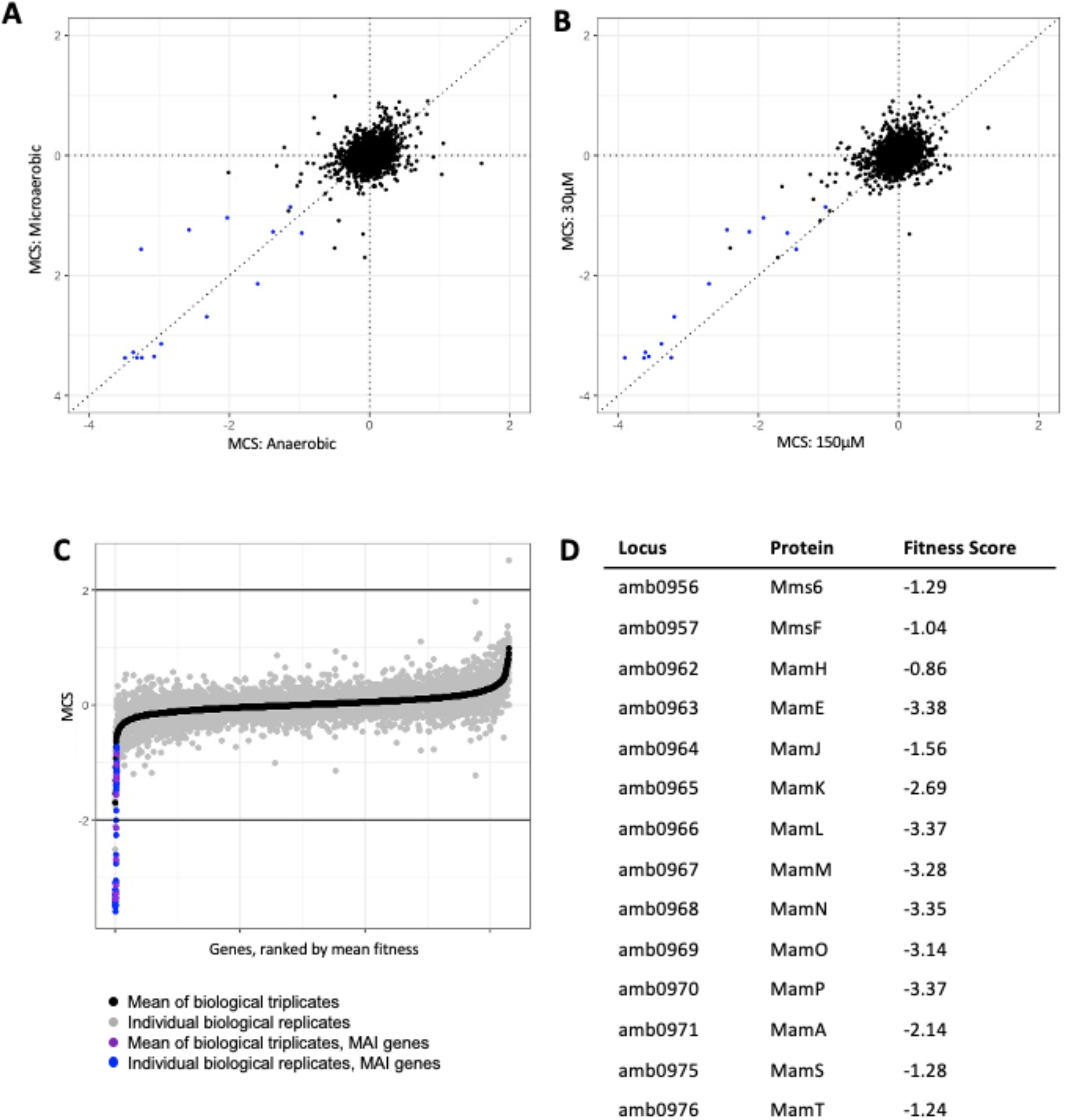
Magnetic fitness of AMB-1 genes (A) Comparison of magnetic fitness scores for AMB-1 grown in microaerobic (<10% oxygen) or anaerobic (0% oxygen) conditions. Each point is the mean gene fitness of three biological replicates. MAI genes with low MCS highlighted in blue (B) Comparison of magnetic fitness scores for AMB-1 grown in 30μM iron or 150μM iron. Each point is the mean gene fitness of three biological replicates. MAI genes with low MCS highlighted in blue (C-D) Magnetosome island gene scores validate the magnetic selection process. (C) Distribution of magnetic fitness scores for WT grown in standard laboratory conditions (microaerobic (<10% oxygen) and 30 μM iron). MAI genes with low MCS highlighted in blue/purple (D) Table highlighting mean fitness scores for MGC genes and their respective proteins.

To validate the screen, ML2 was run over the column after microaerobic growth in 30μM iron. There were too few cells in the non-magnetic sample for preparation of gDNA because there are very few insertions that completely abolish biomineralization. Accordingly, we reasoned that strains with transposon insertions in magnetosome formation genes would be depleted in the magnetic sample. Thus, a low MCS would correspond to a defect in biomineralization (Fig. 3D). Accordingly, we found that strains with transposon insertions in MAI genes with known roles in magnetosome formation—*mms6* (AMB_RS04895), *mmsF* (AMB_RS04900), *mamH* (AMB_RS04920), *mamE* (AMB_RS04930), *mamJ* (AMB_RS23165), *mamK* (AMB_RS04945), *mamL* (AMB_RS04950), *mamM* (AMB_RS04955), *mamN* (AMB_RS04960), *mamO* (AMB_RS04965), *mamP* (AMB_RS04970), *mamA* (AMB_RS04975), *mamS* (AMB_RS04995), and, *mamT* (AMB_RS05000)—were underrepresented in the populations of magnetic cells (Fig. 3D). It is interesting to note that *mamJ* and mamK, which are involved in magnetosome chain alignment, not biomineralization, also had negative MCS under these conditions. Perhaps, uneven chain segregations in these mutants yields a small but detectable subpopulation which cannot bind to the magnetic column.

The goal of further screens was to identify genes involved in biomineralization when cells are grown in alternative conditions. We focused on genes required for biomineralization under anaerobic conditions and under high iron conditions. Recent work by Amor et al. has shown that the concentration of iron supplied in the growth medium affects both iron uptake and magnetite formation in AMB-1 [10]. As iron concentration is increased in the media, cells take up more iron. Additionally, magnetite crystals in AMB-1 cells increase in size as iron concentration is increased. Previous genomic studies have shown that iron uptake genes are upregulated under magnetosome-forming conditions [18, 19, 21]. However, it is not known if there is a genetic response to stimulate increased iron uptake or regulate the size of magnetite crystals in response to iron concentration. Here, we performed a magnetic selection with ML2 after growing cells at standard (30μM) and high (150μM) iron concentrations to uncover genes involved in magnetosome formation when cells are saturated with iron.

After magnetic selection, we compared the MCS from the magnetic populations of the standard and high iron concentrations against each other (Fig. 3B). Most of the MCS either clustered around 0, meaning there was no measurable magnetic defect, or along the slope of 1, meaning there was no difference in magnetic response for that gene between the two iron conditions. There were several genes of interest that were underrepresented (had a magnetic defect) in the magnetic population of each condition. In the standard iron condition, *amb0360* (AMB_RS01830), which is annotated as a hypothetical protein, had a magnetic defect. While there were no protein predictions, *amb0360* does have a homolog in *Magnetospirillum gryphiswaldense* MSR-1 (95.96% identity). One gene, amb4208 (AMB_RS21290) was overrepresented (had a positive MCS) in the high iron condition. It is possible that mutants of this gene are more magnetic than wild-type cells. Again, AMB4208 is a hypothetical protein with homologs in *Magnetospirillum sp*. XM-1 (81.5% identity), ME-1 (80.5% identity), and MSR-1 (75.54% identity).

There were several other genes with magnetic defects in the high iron condition. Of note, there were several MAI genes that were underrepresented in the magnetic population: *mms6, mmsF, mamS*, and *mamT*. MamT is a magnetosome localized protein with a magnetochrome domain (a heme-binding motif unique to magnetosome proteins) and is thought to be involved in redox chemistry for magnetosome crystal growth [26]. MmsF and MamS are also magnetosome-localized proteins and have been shown to regulate crystal size and morphology [11, 12, 27]. The lower magnetic score of these genes in the high iron condition compared to the standard iron condition suggests that they may have a more nuanced function in magnetosome formation than previously thought.

### Magnetic selection reveals genes important for biomineralization under anaerobic conditions

Both AMB-1 and MSR-1 are capable of growing in microaerobic conditions. However, biomineralization only occurs when oxygen concentrations in the medium are depleted [23]. In the laboratory, AMB-1 is typically grown with a low concentration of oxygen (2-10%) in the culture headspace, with nitrate as an alternative electron acceptor. It is not known if the inability of MTB to form magnetosomes aerobically is simply due to the balance of ferrous and ferric iron required to produce magnetosomes or if there is also a genetic response to turn off magnetosome production at high oxygen concentrations.

Here, we grew ML2 in either microaerobic (test tubes with minimal headspace incubated in a microaerobic chamber) or anaerobic (sealed serum bottles with anaerobic media) conditions and performed a magnetic selection to find genes that are required for magnetosome formation in varying oxygen concentrations. The microaerobic conditions reflect how cells are grown in standard lab conditions.

We compared the MCS from the magnetic populations of the microaerobic and anaerobic samples against each other after magnetic selection (Fig. 3A). Again, most of the MCS either clustered around 0 or along the slope of 1, meaning there was no difference in magnetic response for most genes between the two oxygen conditions. In the microaerobic condition, *amb0360*, again had a magnetic defect, suggesting that it may be important for biomineralization under standard conditions. Another gene, *amb4151* (AMB_RS21005), which is annotated as a hypothetical protein, had a positive MCS in the anaerobic condition, suggesting that mutants of this gene are more magnetic than wild-type cells. *Amb4151* has homologs in *Magnetospirillum sp*. XM-1 (89.07% identity), ME-1 (87.89% identity), and MSR-1 (77.66% identity).

Several MAI genes also had magnetic defects in the anaerobic condition: *amb0936* (AMB_RS04810), *amb0947* (AMB_RS04850), *amb1008* (AMB_RS05170), *amb1009* (AMB_RS05175), *amb1022* (AMB_RS23960), *mamT, mmsF*, and *mamJ*. Again, it was surprising that *mamJ* came up in this screen, as MamJ is known for its role in chain organization, not biomineralization. While both *mamT* and *mmsF* deletion strains have defects in biomineralization [26,27], it is surprising that the MCS is even lower in the high iron condition because magnetosomes are typically larger in size under high iron conditions. These results provide further support that the roles of some MAI genes are conditional.

### Δ*mamT*Δ*R9* plays a greater role for biomineralization in high iron and anaerobic conditions

To validate the results of the screen, we chose to look more closely at a *mamT* deletion to confirm that phenotypes seen in the magnetic selection screen also occur when the gene is deleted from a WT background. Here, we used Δ*mamT*Δ*R9*, where region 9 (R9) of the MAI (containing exact duplications of *mamQ,R,B*) was also deleted to prevent a recombination event that makes the *mamT* deletion less stable [26]. When grown with 30μM iron, Δ*mamT*Δ*R9* cells produce smaller crystals and have a very low magnetic response compared to WT cells (Fig. 4A,C). When grown with 150μM iron, TEM images of Δ*mamT*Δ*R9* cells qualitatively look very similar to cells grown with 30μM iron (Fig. 4C). However, the length of the magnetite crystals is decreased in Δ*mamT*Δ*R9* cells in 150μM iron compared to 30μM iron. As described above, it is surprising that magnetosomes would be smaller in the high iron condition. Complementing Δ*mamT*Δ*R9* by integrating *mamT* into the genome restores the WT phenotype in both iron conditions. Again, we validated the phenotype of Δ*mamT*Δ*R9* under anaerobic conditions. When grown anaerobically, Δ*mamT*Δ*R9* magnetite crystals were shorter than both wild-type crystals and Δ*mamT*Δ*R9* under microaerobic conditions (Fig. 4A).

**Figure 4.**
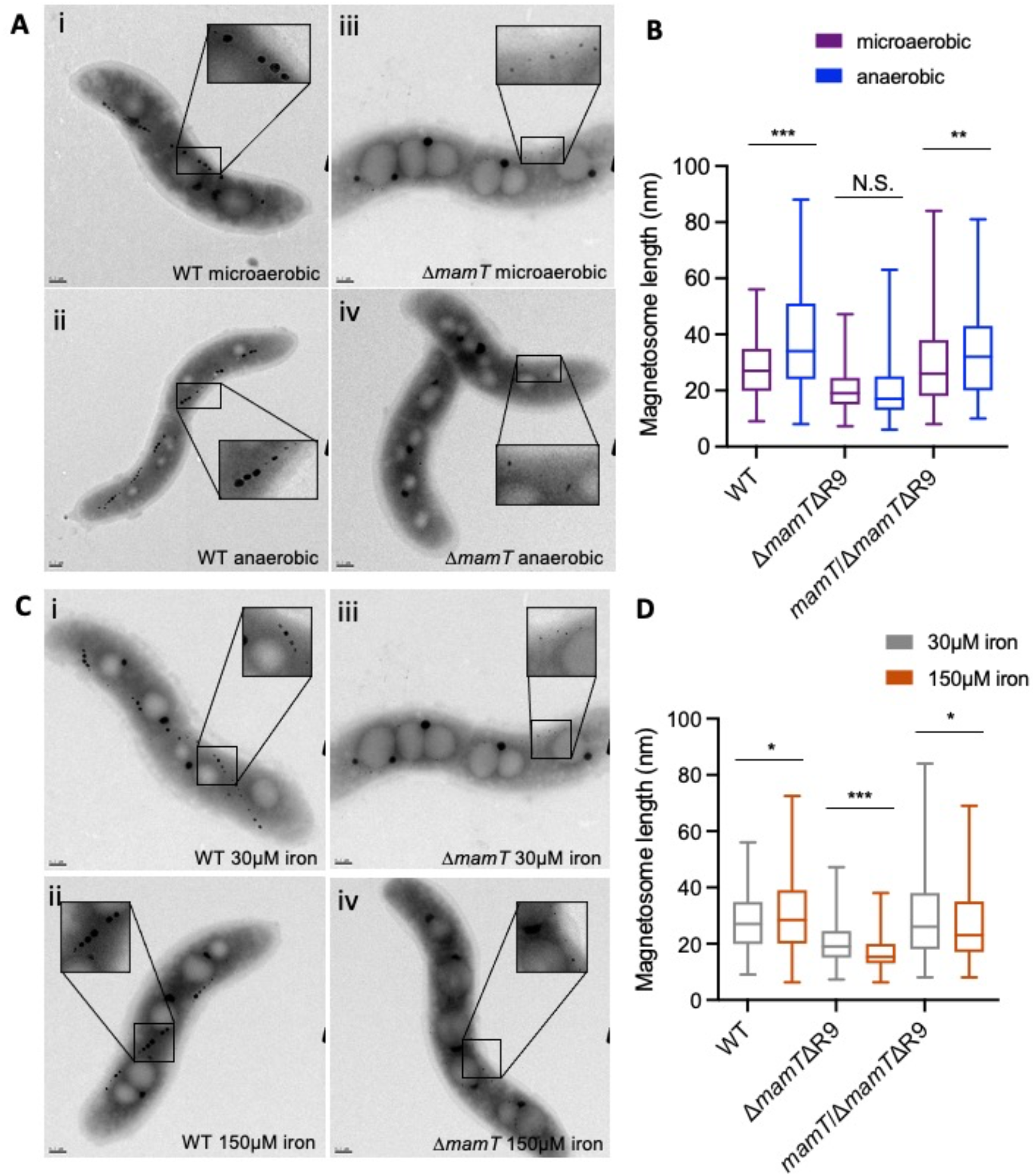
Validation of Δ*mamT* biomineralization phenotypes(A) Representative images of WT and Δ*mamT*Δ*R9* in microaerobic (<10% oxygen) or anaerobic (0% oxygen) conditions. (B) Magnetosome length of WT, Δ*mamT*ΔR9, and *mamT*/Δ*mamT*ΔR9 magnetosomes after growth in microaerobic (<10% oxygen) or anaerobic (0% oxygen) conditions. (C) Representative images of WT and Δ*mamT*Δ*R9* in 30μM iron or 150μM iron conditions. (D) Magnetosome length of WT, Δ*mamT*ΔR9, and *mamT*/Δ*mamT*ΔR9 magnetosomes after growth in 30μM iron or 150μM iron.

All experiments were repeated in the Δ*R9* strain to confirm that phenotypes seen with Δ*mamT*Δ*R9* were not caused by the R9 deletion (Fig. S1). Δ*R9* cells behaved similarly to WT in that they had larger magnetosomes under both high iron and anaerobic conditions. The MAI gene *mms6* had a biomineralization defect in the 30μM iron condition (−1.59 MCS), as expected [28] and slightly less of a defect in the 150μM iron condition (−1.29 MCS), so we looked at Δ*mms6* to verify that the magnetic selection was accurate and as a control for Δ*mamT*Δ*R9*. After growing Δ*mms6* in 30μM or 150μM iron, magnetosomes were larger under the high iron conditions, similar to WT (Fig. S2).

These results together suggest that the magnetic selection process is sufficient for pulling out genes that, when disrupted, have a defect in biomineralization. They also show that *mamT*, which was already known to be involved in magnetosome formation may have a more critical role in both high iron and anaerobic conditions.

### Genes outside the MAI may have a role in biomineralization

It was unexpected to see a gene with a positive MCS in the anaerobic condition, especially a gene outside the MAI, as this could indicate that a novel ex-MAI gene has a role in control of biomineralization. To explore this phenomenon further, we generated a deletion of *amb4151*. Under microaerobic conditions, Δ*amb4151* magnetosomes looked similar to WT magnetosomes (Fig. 5A) and had the same magnetosome length (Fig. 5B). Under anaerobic conditions, WT AMB-1 cells typically produce larger magnetosomes (mean = 37.2 nm, σ = 17.0). Δ*amb4151* cells produced larger magnetosomes than WT under anaerobic conditions and appeared to do so more consistently (mean = 53.5 nm, σ = 15.8).

**Figure 5.**
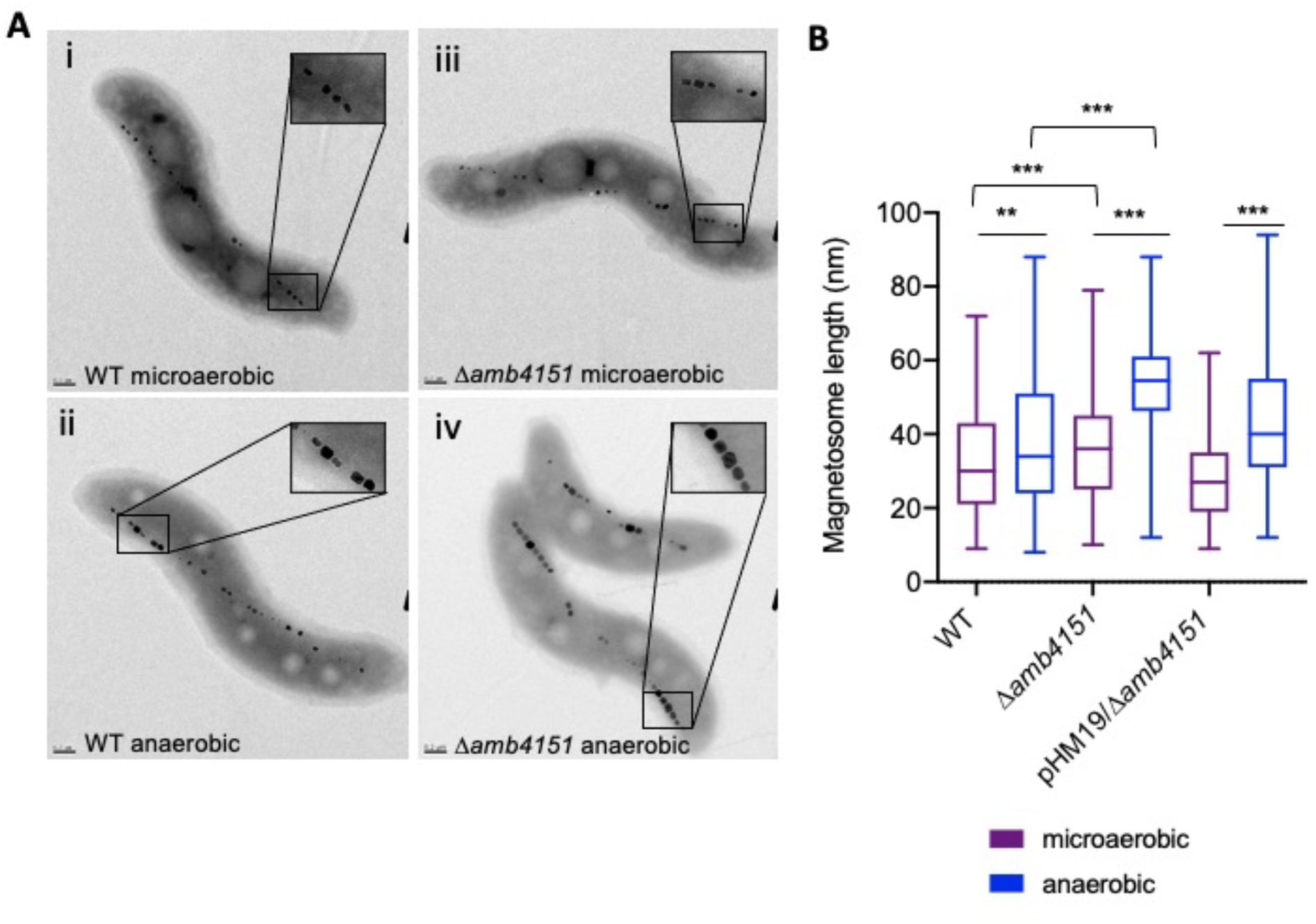
Validation of Δ*amb4151* biomineralization phenotypes. (A) Representative transmission electron micrographs of WT and Δ*amb4151* grown in microaerobic (<10% oxygen) and anaerobic (0% oxygen) conditions. i. WT grown in microaerobic conditions ii. WT grown in anaerobic conditions iii. Δ*amb4151* grown in microaerobic conditions iv. Δ*amb4151* grown in anaerobic conditions (B) Magnetosome length of WT, Δ*amb4151*, and *amb4151*/Δ*amb4151* magnetosomes after growth in microaerobic or anaerobic conditions.

**Figure 6.**
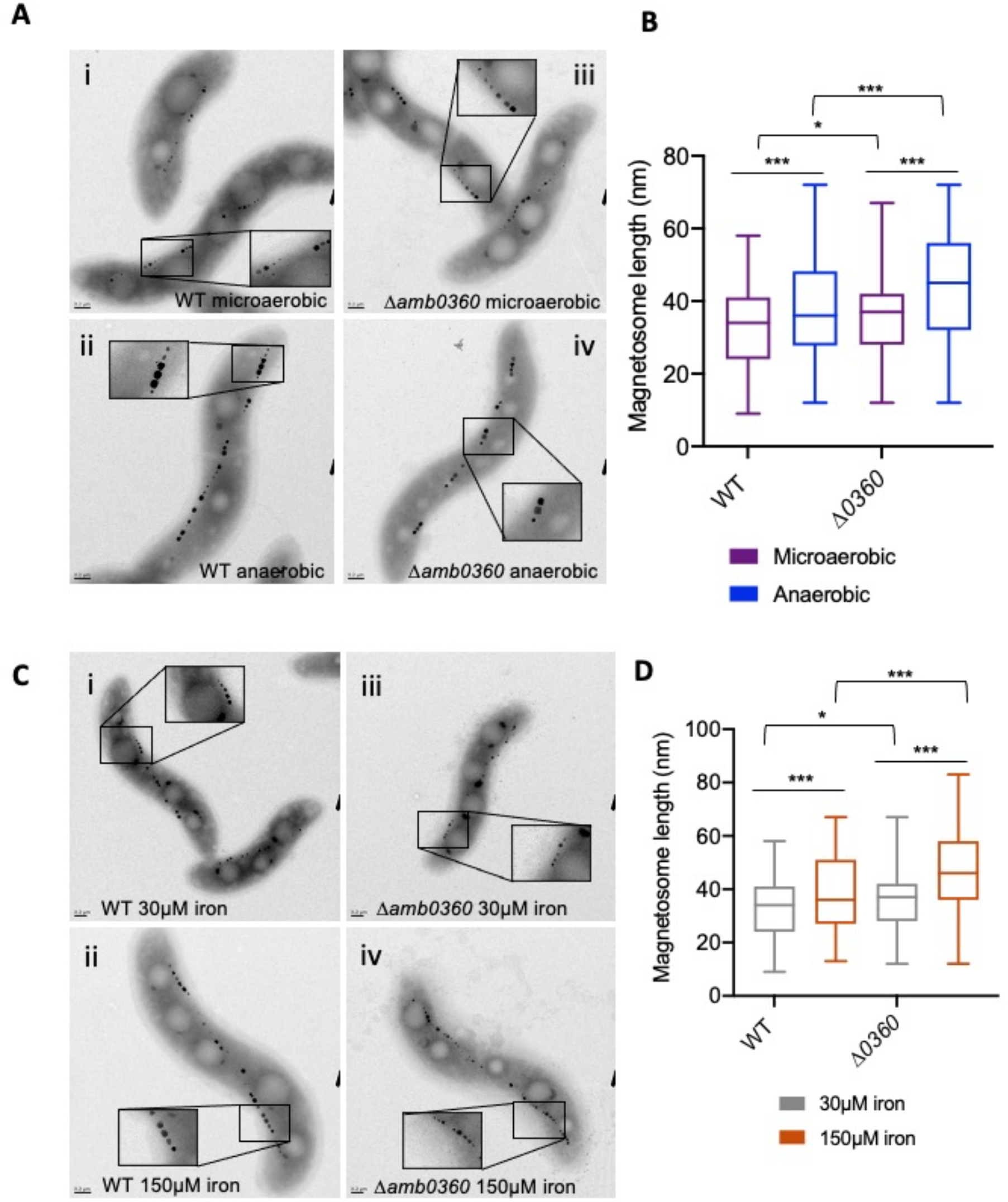
Validation of Δ*amb0360* biomineralization defect. (A) Representative transmission electron micrographs of WT and Δ*amb0360* grown in microaerobic (<10% oxygen) and anaerobic (0% oxygen) conditions. (B) Magnetosome length of WT and Δ*amb0360* magnetosomes after growth in microaerobic or anaerobic conditions. (C) Representative transmission electron micrographs of WT and Δ*amb0360* grown in 30 μM or 150 μM iron. (D) Magnetosome length of WT and Δ*amb0360* magnetosomes after growth in 30 μM or 150 μM iron

## Discussion

Previous genetic studies in MTB have relied on transposon mutagenesis screens that required sorting through thousands of individual colonies [29–31]. Here, we showed that RB-TnSeq is a streamlined and effective technique for investigating biomineralization in MTB, as the pooled library allows for screening hundreds of thousands of mutants through many growth conditions in a relatively short amount of time. Additionally, the method is sensitive to small changes in magnetic properties of a mutant. These features allowed us to demonstrate that some known magnetosome genes have differential roles as growth conditions change and that novel genes outside the MAI may have an impact on biomineralization.

There are clear benefits to using RB-TnSeq over traditional methods of mutagenesis. First, the simplicity of sequencing barcodes in each sample allows for streamlined screening and the ability to test many different conditions in a single experiment. Additionally, because the phenotypic score for a gene is averaged across many individual insertions, we can be more confident that genes of interest are worth pursuing. However, RB-TnSeq is not without its challenges. Because all mutants are pooled together it is possible that some phenotypes of some mutants may be masked via transcomplementation by other mutants in the pool or by the availability of common goods (i.e. siderophores, signaling molecules). Alternatively, it may be difficult to validate the phenotype after screening. There is a chance that a mutant is influenced by competition in the pool—for example, the mutants of one gene are outcompeted for nutrient uptake and form smaller magnetosomes—and once the gene is examined on its own, the phenotype may disappear. Biased insertion and polar effects could also cause false positives in some instances. But a phenotype can still be evaluated after the fact with single gene or operon deletions.

In this study, the ML2 (wild-type) library had full coverage of the genome, with over 180,000 strains and approximately 34 transposon insertions per gene. With ML2 we were able to map the essential gene set of AMB-1 under our standard laboratory conditions, which are the conditions under which most genetic studies have been conducted in AMB-1. The essential gene set expands our understanding of the physiology of AMB-1 and allows for comparison to other MTB in either physiological or phylogenetic studies.

We then screened the transposon mutant pool using a magnetic selection. Previous studies in MTB have used similar selection methods (i.e. colony color, response to a magnetic field, magnetic column) [29–33], but the barcode sequencing adds power by quantifying the results. Since MCS is based on a weighted average of all the mutants in a particular gene, a biomineralization phenotype for a particular gene is more reliable. As the biomineralization phenotype for one strain can vary across the population of cells, averaging the scores of multiple mutants strains provides more confidence that the phenotype is real. Additionally, since ML2 is complete library, this allows us to reexamine mutants that were previously discovered with standard techniques.

Mutants with insertions in MAI genes were depleted from the population of magnetic cells after magnetic selection, indicating that the screening method reliably detects strains with defects in biomineralization. Interestingly, genes identified in a previous transposon mutagenesis screen by Matsunaga et al. (2005) [17] as being essential for biomineralization were not found to have biomineralization defects in this study. Matsunaga et al. created 5762 Tn5 transposon mutants and found 69 mutants with defects in biomineralization, none of which were located in the MAI. It is possible there were secondary mutations in the Tn5 transposon mutants that were the true cause of the biomineralization defect—for example, spontaneous loss of the MAI. One benefit of RB-TnSeq is that the phenotype is based on an average of multiple insertions in each gene, making it less likely to falsely link a gene to biomineralization.

We also discovered potential new functions for known magnetosome genes when cells were grown under non-standard conditions (high iron or anaerobic). Of note was *mamT* which is known to be involved in crystal formation but may be more critical under high iron or anaerobic conditions. The *mamT* phenotype is notable because under high iron or anaerobic conditions we expect magnetosomes to get larger, and *mamT* mutants have the opposite phenotype. The effect on magnetosome size was also quite subtle, which highlights the power of the magnetic screen and is ability to identify phenotypes that would have been overlooked otherwise. While we did not investigate the mechanism in this work, perhaps the decreased size of magnetosomes in those two conditions are due to the proposed role of MamT in regulating the balance of iron species within the magnetosome. Further study of MamT under alternative growth conditions could provide further insight into its function.

While some of the genes in the MAI are transposable elements and are unlikely to have a role in magnetosome formation, the screens here showed that many of the genes in the MAI originally thought to have no role in biomineralization—or that have not been studied before—may be needed as conditions shift. These genes include: *amb0936, amb0947, amb0958, amb0959, amb1009, amb1008, amb1022, mamJ, mamK*, and *mamY*. It is especially surprising that *mamJ* mutants had a biomineralization defect, since the phenotype of *mamJ* is masked by the presence of *limJ* in the magnetosome islet [34]. Additionally, we uncovered genes outside the MAI that may have an impact on magnetosome formation. We found that *amb4151*—annotated as a hypothetical protein—inhibits biomineralization to a small degree. Δ*amb4151* has larger magnetosomes under anaerobic conditions, when compared to WT.

It has been shown that magnetosomes are only made under low oxygen concentrations [23]. However, it is not known if this is simply due to shifting redox conditions or if there is also a genetic response to turn off magnetosome production at high oxygen concentrations. Further study of *amb4151* could be useful to understand the impact of ex-MAI genes on magnetosome formation. The phenotype of *amb4151* also suggests that the changes in magnetosome size under shifting iron and oxygen increases are not only due to redox conditions but are also influenced by a genetic response.

The genes highlighted in this study all had nuanced phenotypes, in that each gene deletion had a slight effect on biomineralization. While these genes may not be critical to the formation of magnetosomes, it is still valuable to study their functions. Most, if not all, of the genes that have a large impact on biomineralization or magnetosome formation have already been identified in past screens and reverse genetic studies. Genes with more subtle phenotypes can provide key information on the formation of magnetosomes under different conditions.

While previous studies have suggested that MTB may have a large environmental impact, we know very little about the dynamics of magnetosome formation in natural environments. A greater understanding of the molecular mechanisms of biomineralization can help with understanding how much iron is taken up by MTB and under what conditions. This study shows that RB-TnSeq and screens for biomineralization defects are useful tools to identify genes that influence the response of AMB-1 to changing environmental conditions. There are certainly many applications for this technique to reveal more about growth and biomineralization in MTB. For example, an RB-TnSeq library may be used to find genes that are necessary in an oxygen or redox gradient. Or taken a step further, could be used to find mutants in magnetoaerotaxis. Both screens would reveal more about the physiology and lifestyle of magnetotactic bacteria.

## Methods

### Growth and culture conditions

*Magnetospirillum magneticum* AMB-1 was cultured in defined minimal media (MG medium,) supplemented with 1/100 vol of Wolfe’s vitamin solution and 30μM ferric malate as previously described [31]. Colonies were grown on solid MG with 0.7% agar. Kanamycin was used at 10μg/mL in solid media and 7μg/mL in liquid media for strains with a kanamycin-resistant cassette integrated into the chromosome or on a plasmid. For microaerobic growth cells were grown in culture tubes or 50mL conical tubes and incubated at 30°C in a microaerobic chamber with 10% oxygen. For anaerobic growth, sealed Balch tubes containing 10mL of MG medium and 20mL of headspace were used. Media was bubbled with N_2_ gas for 10 min before sealing. Then headspace was flushed with N_2_ gas for 10 min before autoclaving. Ferric malate and Wolfe’s vitamins were added once tubes cooled. All cultures inoculated with a dilution factor of 1:100.

### Library construction + processing

Detailed description of transposon vector construction is described in Wetmore et al., 2015 [22]. AMB-1 mutant libraries were constructed using APA752 (*mariner* transposon library pKMW3 in WM3064 containing millions of unique 20-nucleotide barcodes). APA752 was transferred to AMB-1 from WM3064 by conjugation: 1L of wild-type AMB-1 culture at OD400 0.230 was conjugated with 100mL of APA752 culture by incubating in a microaerobic (10% oxygen) chamber for 14 hours, then diluted 1:15 and plated on MG/Kan. Plates were incubated in microaerobic jars (7% oxygen) at 30°C for 5 days. Colonies were collected from plates and pooled in liquid MG/Kan, then allowed to grow to late log phase before spinning down 50 mL aliquots and freezing in 1 mL aliquots with 20% glycerol. All libraries were made in standard MG. Some cell pellets from the 50 mL aliquots were set aside for gDNA extraction. To ensure that the library had enough barcodes to indicate complete coverage of the genome, BarSeq PCR followed by Illumina MiSeq sequencing was performed. Libraries determined to be diverse enough were sequenced by Illumina HiSeq to map transposon insertions. Illumina library preparation and TnSeq data analysis are described in Wetmore et al., 2015 [22].

### Essential gene analysis

We used previously published methods to determine the essential gene set of AMB-1 in both standard wild-type and Δ*MAI* strains using the RB-TnSeq libraries [25]. Briefly, read density was determined for each protein-coding gene. Genes that are very similar to other parts of the genome and genes of less than 100 nucleotides were excluded from analysis. Based on the median insertion density and median length of remaining genes, a threshold for gene size included in the essential gene analysis was set for the library by determining how short a gene could be and still be unlikely to have no insertions at all by chance (*P*<0.02, Poisson distribution) [25]. The threshold for ML2 was 800 nucleotides—any genes shorter than that were excluded from the essential gene analysis.

### Competitive mutant magnetic assays

For each condition, one aliquot of ML2 was thawed and inoculated into 250mL of MG with 7 μg/mL Kanamycin. When the culture recovered to stationary phase—~OD400 0.250 on Ultrospec 2100 pro (Amersham)—cell pellets were collected from 25mL aliquots as time 0 samples. For experiments testing iron concentrations, cells were first washed twice and resuspended in MG without carbon or iron, respectively. Then, cultures were inoculated at 1:100, in triplicate for each condition. For iron experiments, MG was supplemented to either 30μM or 150μM ferric iron using 3mM ferric malate stock. Cultures were balanced with malic acid (18mM (200x) stock). For oxygen experiments with ML2, microaerobic cultures were grown in culture tubes in a microaerobic glove box without shaking and anaerobic cultures were grown in Balch tubes without shaking.

### Magnetic selection

15mL of culture from each condition and each triplicate were pelleted as the pre-column sample and frozen at −20°C. For each magnetic selection, an LS column (Miltenyi Biotech) was set up on a ring stand with 6 sets of 2 disc magnets on each side. The remaining 35mL of culture were passaged over the magnetic column via gravity filtration and collected in a 50mL conical (magnetic sample). Then, magnets were removed and column was washed twice with 5mL of MG. Flow-through (non-magnetic sample) was collected. Both magnetic and non-magnetic samples were spun down and cell pellets were frozen at −20°C.

### BarSeq

Genomic DNA was isolated from library samples using the Qiagen DNeasy Blood & Tissue Kit. gDNA concentration was quantified by NanoDrop. BarSeq PCR was performed using ~200ng of DNA for each sample. Barcodes were amplified using unique, indexed primers, as described in Liu et al., 2021 [35]. Samples were pooled and sequenced using Illumina HiSeq single-end reads.

### BarSeq data analysis and magnetic column abundance calculation

A detailed description of BarSeq analysis can be found in Wetmore et al., 2015. Briefly, the magnetic column abundance score (MCS) for a gene is calculated from the weighted average of strain MCS. BarSeq data can be viewed in the Fitness Browser (http://fit.genomics.lbl.gov), which provides information from successful selection experiments, including details of the experimental conditions, quality metrics for each experiment, per-strain scores, and gene scores [25]. Because these methods were developed for use with growth assays, we added an additional normalization step to account for any growth defects that might influence MCS after passing cells through the magnetic column by subtracting pre-column gene MCS from magnetic and non-magnetic gene MCS.

### Strains

Δ*mamT*Δ*R9* [26], Δ*mmsF* [27], and Δ*mms6* [27] were generated previously. All other gene deletions were made using a two-step recombination method to generate a markerless deletion [31]. Deletion plasmids were constructed using three-piece Gibson assembly. Regions upstream and downstream of the gene of interest were cloned into pAK31 (suicide vector containing the Kanamycin-resistant cassette and *sacB* gene) cut with BamH1 and Spe1. The plasmids were transferred to AMB-1 cells by conjugation and selected on MG/Kanamycin media. Colonies were then grown on MG with 2% sucrose to select for deletion strains. Sucrose-resistant colonies were screened by PCR for the deletion and for the lack of plasmid markers.

Complementation plasmids were constructed using three-piece Gibson assembly. The gene of interest was cloned into pAK22 (replicative plasmid containing the Kanamycin-resistant cassette following a tac promoter) cut with EcoR1 and Spe1. The plasmids were transferred to AMB-1 cells by conjugation and selected on MG/Kanamycin media.

### Plasmids

**Plasmids:**
| Name | Origin | Description | Reference |
| --- | --- | --- | --- |
| pHM14 | pAK31-derived | Deletion plasmid for *amb4151* | This work |
| pHM19 | pAK22-derived | Complementation plasmid for $\Delta amb4151$ | This work |
| pAK807 | pAK605-derived | Complementation plasmid for $\Delta mamT\Delta R9$ | Patrick Browne (unpublished) |
| pHM20 | pAK31-derived | Deletion plasmid for *amb3952* | This work |

### Strains

**Strains:**
| Strain | Organism | Description | Reference |
| --- | --- | --- | --- |
| APA752 | *E. coli* | mariner RB-TnSeq library | Wetmore et al. |
| AK155 | AMB-1 | $\Delta mamT\Delta R9$ | Jones et al. |
| AK57 | AMB-1 | $\Delta R9$ | Murat et al., 2010 |
| AK104 | AMB-1 | $\Delta mmsF$ | Murat et al., 2012 [27] |
| AK110 | AMB-1 | $\Delta mms6$ | Wan et al. (in review) |
| HM02 | AMB-1 | *Δamb4151* | This work |
| DH5α (λpir) | *E. coli* | Standard molecular cloning strain |  |
| WM3064 | *E. coli* | Plasmid conjugation strain |  |

### Primers

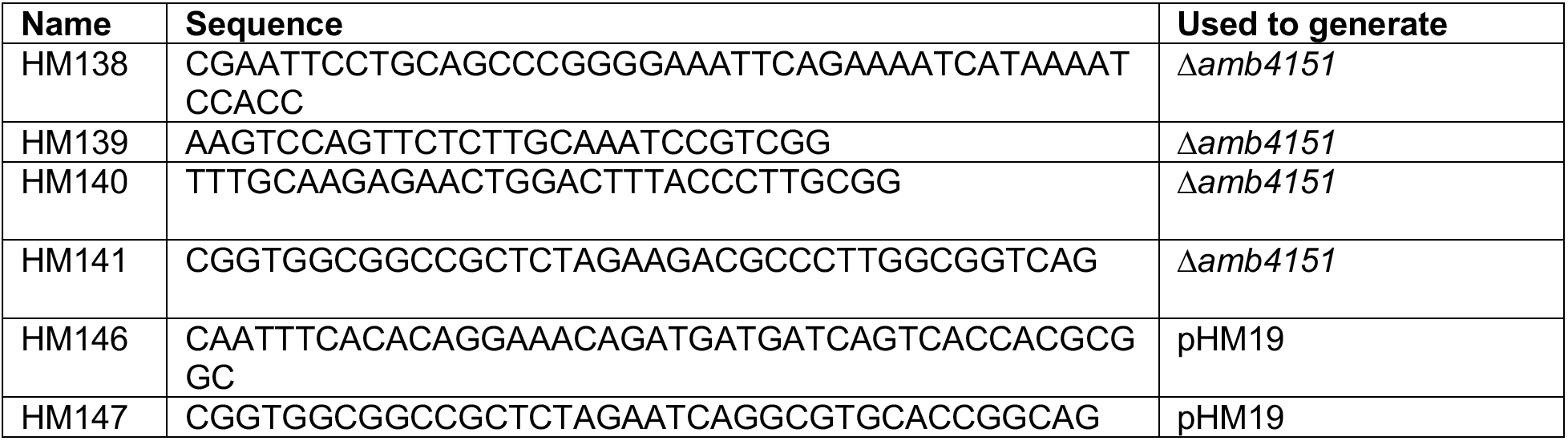

### TEM

Whole AMB-1 cells were imaged by transmission electron microscopy (TEM). To prepare samples for imaging, 1.5mL of stationary phase (~OD_400_ 0.25) culture was pelleted and resuspended in 10μL of MG. Cells were absorbed on glow-discharged, 200-mesh Cu grid with Formvar film and imaged on FEI Technai 12 transmission electron microscope with a Gatan Bioscan (1,000 by 1,000) charge-coupled device (CCD) camera model 792 at an accelerating voltage of 120kV.

### Quantification of crystal size and magnetosome numbers

TEM images were used to measure crystal size and numbers. The length and width of each crystal was measured along its long axis by hand using ImageJ.

## Supporting information

Magnetic Selection Fitness Score

## Acknowledgements

We thank Morgan Price for assisting with the analysis of RB-TnSeq and BarSeq data and faculty at the EM-Lab at the University of California, Berkeley for their assistance with TEM. This material by ENIGMA-Ecosystems and Networks Integrated with Genes and Molecular Assemblies (http://enigma.lbl.gov), a Science Focus Area Program at Lawrence Berkeley National Laboratory is based upon work supported by the U.S. Department of Energy, Office of Science, Office of Biological & Environmental Research under contract number DE-AC02-05CH11231. This work was funded by generous support from the NIH (R35GM127114) and the Miller Institute for Basic Research in Science. Hayley McCausland’s work was partially supported by the Molecular Basis of Cell Function Training Grant (T32GM007232).

## Supplemental materials

**Figure S1.**
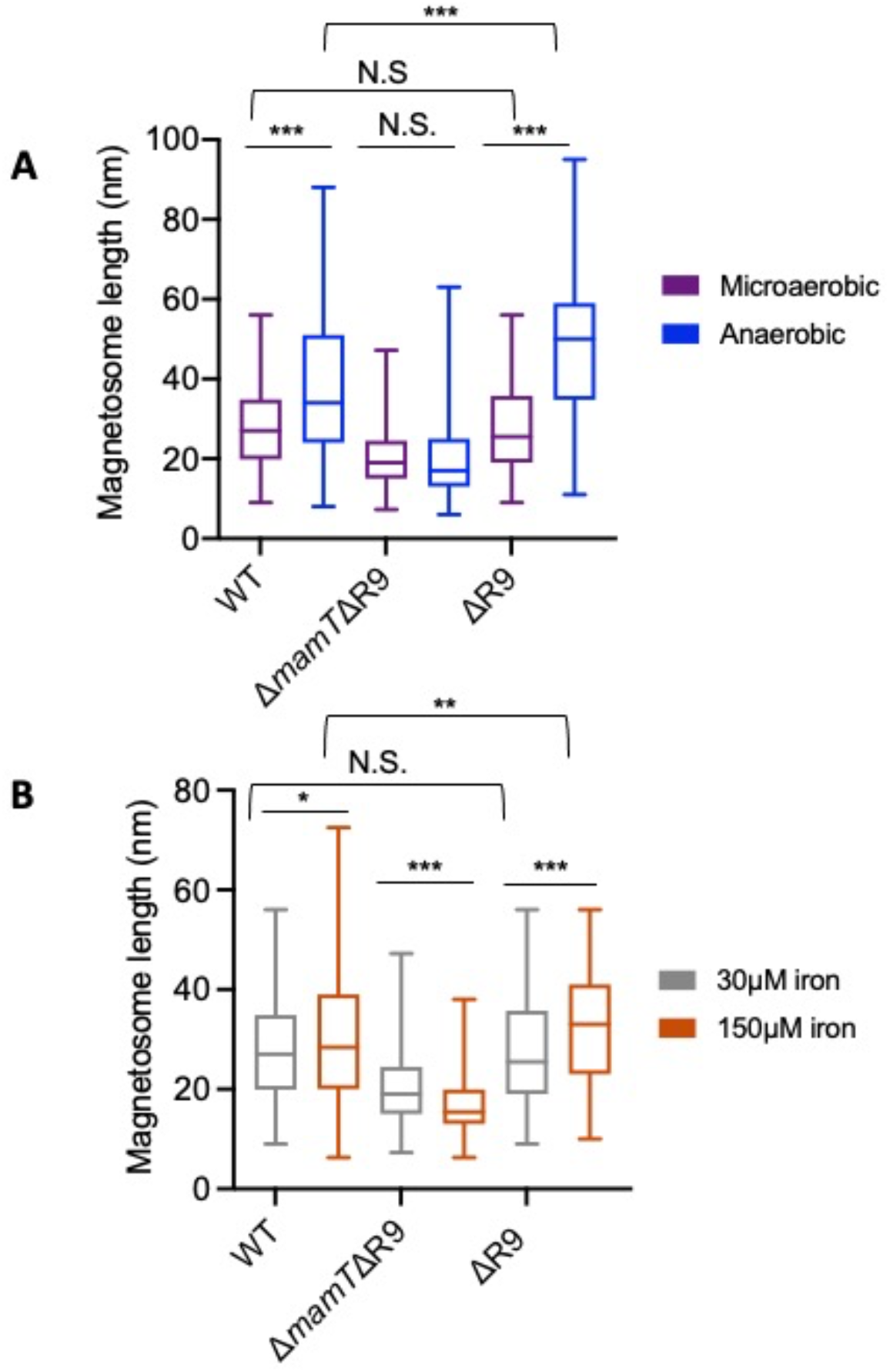
Δ*R9* does not impact the Δ*mamT*Δ*R9* phenotype (A) Magnetosome length of WT, Δ*mamT*Δ*R9*, and Δ*R9* magnetosomes after growth in microaerobic (<10% oxygen) or anaerobic (0% oxygen) conditions. (B) Magnetosome length of WT, Δ*mamT*Δ*R9*, and Δ*R9* magnetosomes after growth in 30 μM iron or 150 μM iron conditions.

**Figure S2.**
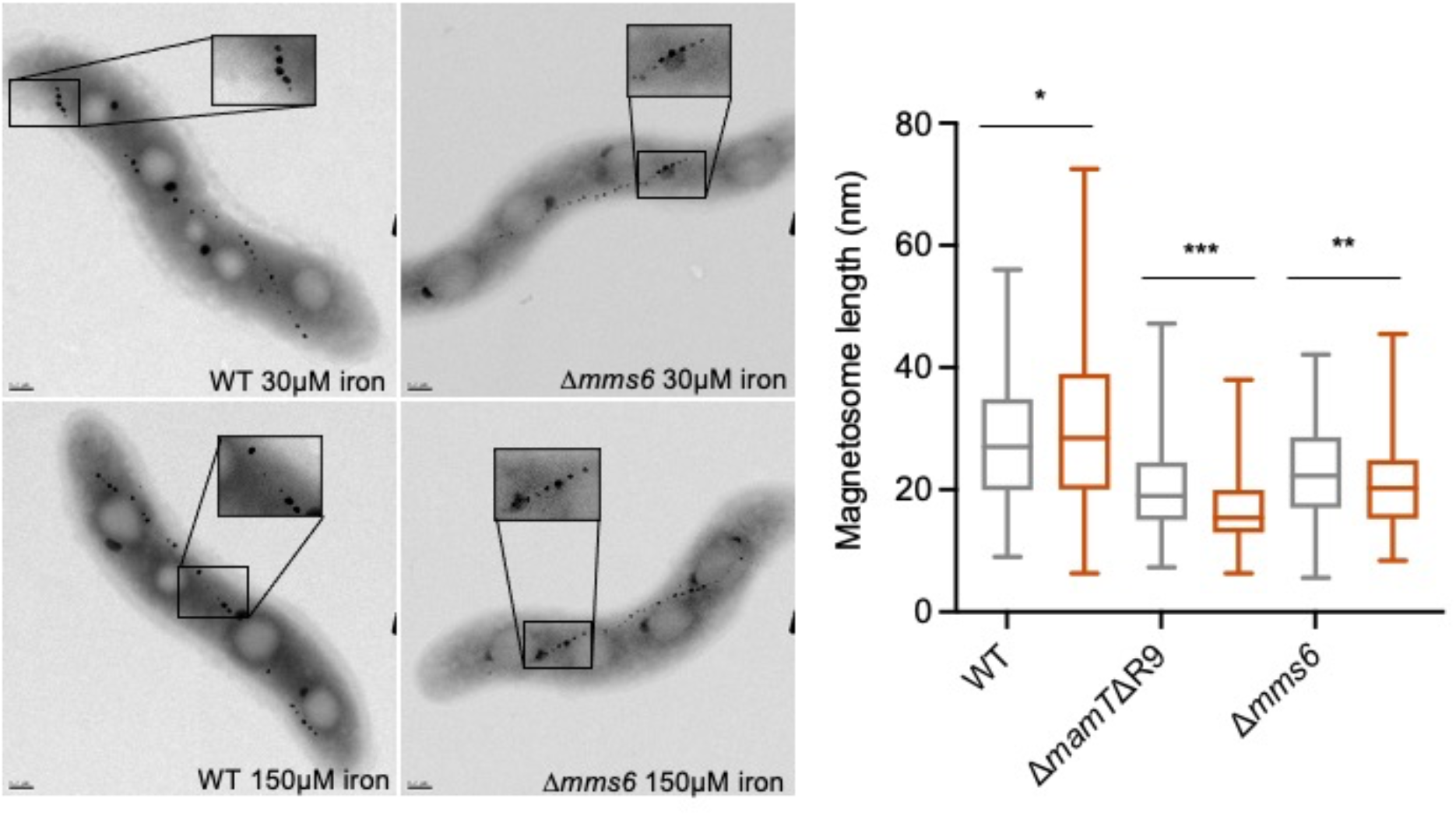
Δ*mms6* magnetosomes are longer under high iron conditions. (A) Representative transmission electron micrographs of WT and Δ*mms6* grown in 30 μM or 150 μM iron. (B) Magnetosome length of WT, Δ*mamT*Δ*R9*, and Δ*mms6* magnetosomes after growth in 30 μM iron or 150 μM iron conditions.

**Table S1.** Test for normality in magnetosome length analysis

| Figure | Strain, Condition | Shapiro-Wilk normality test (P-value) |
| --- | --- | --- |
| Fig. 4B | WT, microaerobic | 0.0054 |
|  | WT, anaerobic | <0.0001 |
| | $\Delta mamT\Delta R9$, microaerobic | <0.0001 |
| | $\Delta mamT\Delta R9$, anaerobic | <0.0001 |
| | $mamT/\Delta mamT\Delta R9$, microaerobic | <0.0001 |
| | $mamT/\Delta mamT\Delta R9$, anaerobic | <0.0001 |
| Fig. 4D | WT, 30 μM | 0.0054 |
| | WT, 150 μM | <0.0001 |
| | $\Delta mamT\Delta R9$, 30 μM | <0.0001 |
| | $\Delta mamT\Delta R9$, 150 μM | <0.0001 |
| | $mamT/\Delta mamT\Delta R9$, 30 μM | <0.0001 |
| | $mamT/\Delta mamT\Delta R9$, 150 μM | <0.0001 |
| Fig. 5B | WT, microaerobic | <0.0001 |
|  | WT, anaerobic | <0.0001 |
| | $\Delta amb4151$, microaerobic | 0.002 |
| | $\Delta amb4151$, anaerobic | <0.0001 |
| | pHM19/$\Delta amb4151$, microaerobic | <0.0001 |
| | pHM19/$\Delta amb4151$, anaerobic | <0.0001 |
| Fig. S1A | WT, microaerobic | 0.0054 |
|  | WT, anaerobic | <0.0001 |
| | $\Delta mamT\Delta R9$, microaerobic | <0.0001 |
| | $\Delta mamT\Delta R9$, anaerobic | <0.0001 |
| | $\Delta R9$, microaerobic | <0.0001 |
| | $\Delta R9$, anaerobic | 0.0238 |
| Fig. S1B | WT, 30 μM | 0.0054 |
| | WT, 150 μM | <0.0001 |
| | $\Delta mamT\Delta R9$, 30 μM | <0.0001 |
| | $\Delta mamT\Delta R9$, 150 μM | <0.0001 |
| | $\Delta R9$, 30 μM | <0.0001 |
| | $\Delta R9$, 150 μM | 0.0003 |

**Table S2.** Significance difference tests between magnetosome length datasets

| Figure | Strains/Conditions | Mann-Whitney U test |  | t-test |  |
| --- | --- | --- | --- | --- | --- |
|  |  | p-value | Significant difference | p-value | Significant difference |
| Fig. 4B | WT microaerobic & WT anaerobic | 0.030 | * | ---- | ---- |
| | $\Delta mamT\Delta R9$ microaerobic & $\Delta mamT\Delta R9$ anaerobic | 0.061 | * | ---- | ---- |
| | $mamT/\Delta mamT\Delta R9$ microaerobic & $mamT/\Delta mamT\Delta R9$ anaerobic | 0.007 | ** | ---- | ---- |
| | WT microaerobic & $\Delta mamT\Delta R9$ microaerobic | <0.0001 | **** | ---- | ---- |
| | WT anaerobic & $\Delta mamT\Delta R9$ anaerobic | <0.0001 | **** | ---- | ---- |
| | WT microaerobic & $mamT/\Delta mamT\Delta R9$ microaerobic | 0.580 | N.S. | ---- | ---- |
| | WT anaerobic & $mamT/\Delta mamT\Delta R9$ anaerobic | 0.023 | * | ---- | ---- |
| Fig. 4D | WT 30 μM & WT 150 μM | 0.030 | * | ---- | ---- |
| | $\Delta mamT\Delta R9$ 30 μM & $\Delta mamT\Delta R9$ 150 μM | <0.0001 | **** | ---- | ---- |
| | $mamT/\Delta mamT\Delta R9$ 30 μM & $mamT/\Delta mamT\Delta R9$ 150 μM | 0.041 | * | ---- | ---- |
| | WT 30 μM & $\Delta mamT\Delta R9$ 30 μM | <0.0001 | **** | ---- | ---- |
| | WT 150 μM & $\Delta mamT\Delta R9$ 150 μM | <0.0001 | **** | ---- | ---- |
| | WT 30 μM & $mamT/\Delta mamT\Delta R9$ 30 μM | 0.580 | N.S. | ---- | ---- |
| | WT 150 μM & $mamT/\Delta mamT\Delta R9$ 150 μM | 0.0004 | *** | ---- | ---- |
| Fig. 5B | WT microaerobic & WT anaerobic | 0.0015 | ** | ---- | ---- |
| | $\Delta amb4151$ microaerobic & $\Delta amb4151$ anaerobic | <0.0001 | **** | ---- | ---- |
| | pHM19/$\Delta amb4151$ microaerobic & pHM19/$\Delta amb4151$ anaerobic | <0.0001 | **** | ---- | ---- |
| | WT microaerobic & $\Delta amb4151$ microaerobic | 0.007 | ** | ---- | ---- |
| | WT anaerobic & $\Delta amb4151$ anaerobic | <0.0001 | **** | ---- | ---- |
| | WT microaerobic & pHM19/$\Delta amb4151$ microaerobic | 0.003 | ** | ---- | ---- |
| | WT anaerobic & pHM19/$\Delta amb4151$ anaerobic | <0.0001 | **** | ---- | ---- |
| Fig. S1A | $\Delta R9$ microaerobic & $\Delta R9$ anaerobic | <0.0001 | **** | ---- | ---- |
| | WT microaerobic & $\Delta R9$ microaerobic | 0.828 | N.S. | ---- | ---- |
| | WT anaerobic & $\Delta R9$ anaerobic | <0.0001 | **** | ---- | ---- |
| Fig. S1B | $\Delta R9$ 30 μM & $\Delta R9$ 150 μM | <0.0001 | **** | ---- | ---- |
| | WT 30 μM & $\Delta R9$ 30 μM | 0.828 | N.S. | ---- | ---- |
| | WT 150 μM & $\Delta R9$ 150 μM | 0.008 | ** | ---- | ---- |

Significantly different (\* $P < 0.05$ , \*\* $P < 10^{-2}$ , \*\*\* $P < 10^{-3}$ , \*\*\*\* $P < 10^{-4}$ ); N.S. not significantly different.

## Notes

### Competing Interest Statement

The authors have declared no competing interest.

